# Joint effect of temperature and insect chitosan on the heat resistance of *Bacillus cereus* spores in rice derivatives

**DOI:** 10.1101/2022.04.28.489901

**Authors:** Maria Ines Valdez, Ubeda Maria, Narvaes Cristian, Rodrigo Dolores, Martinez Antonio

## Abstract

The heat resistance of *Bacillus cereus* spores inoculated in a rice substrate supplemented with insect chitosan as an alternative antimicrobial was studied. Two concentrations of insect chitosan were considered in order to assess the role of the insect chitosan concentration during the heat process as an antimicrobial replacement for crustacean chitosan.

Results of the study indicated that the D_T_ values were clearly higher in the substrate without chitosan than in the substrate containing chitosan thus indicating a greater heat resistance to heat treatment of the microorganism inoculated in the substrate without chitosan. This behaviour was also evidenced in the survival curves. There were no great differences between either of the insect chitosan concentrations tested regarding the D_T_ values. The z values were 9.8 ºC on rice substrate. 8.9 ºC on rice substrate supplemented with insect chitosan at 150 µg/mL and 10.7 ºC on rice substrate supplemented with 250 µg/mL of insect chitosan, the chitosan concentration appears to affect the z value of the microorganism. Our results indicate that the combination of heat with insect chitosan as an antimicrobial on foodstuffs subjected to cooking is feasible and can improve the safety of rice derivatives.

## Introduction

*Bacillus cereus* is present in many foods due to its ubiquitous nature. This microorganism is one of the top ten pathogens responsible for many foodborne diseases in humans (Rodrigo et al.. 2021). According to the latest EFSA and ECDC report (EFSA and ECDC 2021) *B. cereus* was involved in 38 outbreaks of strong evidence and 117 outbreaks of weak evidence with a total of 155 outbreaks reported in 2019. Some recent outbreaks in non-EU countries have also been associated with this pathogen; 45 people were affected in an outbreak in a restaurant in Canberra (Australia) in 2018 (Thirkell et al.. 2019) and 200 students in an outbreak in a school in China in 2018 (Chen et al.. 2019).

*Bacillus cereus* causes two types of food poisoning one of an emetic nature and the other of a diarrheal nature (Griffiths and Schraft. 2017). On the one hand diarrheal syndrome is caused by a gastrointestinal disorder due to the ingestion of *B. cereus* spores present in food and at a dose given, an appreciable probability that dells cross the stomach barrier and implanting themselves in the small intestine is possible. Once they germinate in the small intestine they produce enterotoxins that cause disease. On the other hand emetic syndrome is associated with the production of cereulide toxin in the food contaminated with spores that germinate and produce the toxin resulting in foodborne poisoning (Rouzeau et al.. 2020).

In general, this microorganism is associated with complex food products that may include rice as a component; however, other rice-based products and farinaceous foods such as pasta and noodles are also frequently contaminated and involved in cases of *B. cereus* poisoning (Grande et al.. 2006).

The ability of *B. cereus* to form spores and biofilms enables its persistence in various ecological niches and food products resulting in its presence in processed foods such as cooked rice (Navaneethan and Effarizah. 2021). Furthermore, it is the bacteria most commonly present in rice and rice-based products (Hwang and Huang. 2019).

Rice is a basic cereal in many diets and is widely consumed by the general population given its ample supply of nutrients and its relatively low cost. This cereal is one of the most important staple crops feeding almost half of the world’s population (Weıand Huangm 2019). Starch is the most abundant component of a rice grain constituting about 80% of the dry weight of a brown rice grain and approximately 90% of a milled rice grain (Bao 2019). Rice also provides an important variety of micronutrients including vitamins such as niacin thiamine, pyridoxine or vitamin E. and minerals such as potassium, phosphorus, magnesium and calcium (Base de Datos Española de Composition de Alimentos 2021). These conditions provide a very good substrate for *B. cereus* growth and subsequent toxin production.

This cereal is habitually contaminated by *B. cereus* spores throughout all production stages from cultivation to the later stages of processing and consumption. It is believed that the primary habitat of emetic strains could be related to roots tubers and mycorrhizae of some plants such as rice which could explain the generally higher prevalence of these strains in carbohydrate-rich foods. In fact, starch has been shown to promote *B. cereus* growth and emetic toxin production. This would explain why most outbreaks of emetic disease are associated with starch-rich farinaceous foods (Ehling-Schulz et al.. 2015).

Some works pointed out that the current cooking processes for rice and rice derivatives do not inactivate *B. cereus* spores and consequently they can germinate and grow in food if it is not stored properly (Rodrigo et al. 2021). Different control measures have been proposed to control *Bacillus cereus* in foods. As an additional strategy, heat treatment can be combined with other control measures. In this respect, crustacean chitosan has received attention as antimicrobial. It is a polysaccharide with a well-documented antibacterial activity towards vegetative cells which has already been effectively applied as edible chitosan films (Elsabee 2014) and in food packaging applications (Kumar et al… 2020; Priyadarshi and Rhim. 2020). Nevertheless insect chitosan no currently used as antimicrobial could be an alternative to the crustacean chitosan as an additional control measure applied during heat processing of rice thus favouring the destruction of *B. cereus* spores by affecting their heat resistance. According to Van Huis *et al*. (2013) rearing insects is a sustainable activity more friendlily with the environment than fishing or traditional farming. Currently there are no data on the joint effect of insect chitosan and heat on the heat resistance of *B. cereus* spores since chitosan from crustaceans is used as a natural antimicrobial in the preservation processes.

The purpose of this study is to determine how *B. cereus* spore inactivation is affected by the presence of insect chitosan during the heat treatment. This knowledge can pave the way to a better control of *B. cereus* during and after the cooking processes of rice and its derivatives.

## Material and methods

### Microorganisms and sporulation procedure

The *Bacillus cereus* CECT 148 strain used in this study was obtained from the Spanish Type Culture Collection (CECT), (Valencia, Spain). The strain was reactivated in nutrient broth by shaking for 24 hours at 32 ºC and subsequently 0.5 mL of the *B. cereus* culture was inoculated in 20 Roux flasks (Fisher Scientific SL, Madrid, Spain) with Fortified Nutritive Agar (Scharlab. Barcelona, Spain) and incubated at 30 ºC. When the sporulation level reached approximately 90% the spores were collected.

Spore harvesting was performed using a modified metal Digralsky loop (Deltalab, Barcelona, Spain) gently sweeping the agar surface and washing it with double distilled water. The collected solution was centrifuged at 2500g for 15 minutes at 5ºC the supernatant was removed suspended again in 5mL of double distilled water and was centrifuged under the same previously described conditions this process was repeated 4 times. Finally, the spores from the pellet were stored at 4 ºC in distilled water.

### Substrate preparation

The rice solutions (cooked and lyophilized rice) were prepared by dissolving 0.4 g in 19 mL H_2_O. All solutions were sterilized by filtration through of a 0.45µm filter. After sterilizing the rice solution, 1 mL of the spore solution was added and homogenous distribution was guaranteed by a vortex.

Two solutions of rice with chitosan (150 and 250 µg/mL chitosan) (ecoProten, Cordoba, Spain) were used for the heat resistance studies on the food matrix. The pH was adjusted to between 6.8 and 6.9 by using NaOH. Finally, 1 mL of the spore suspension was added and homogenous distribution was guaranteed by a vortex. The resulting 20 mL of solution containing spores and chitosan were poured into a 50 mL sterile beaker.

In all cases the spore concentration in the resulting rice solution was 108 spores/mL.

### Capillary filling and heat treatment

The capillary tubes with one end closed were supplied by Vitrex, reference 217913 (1.50 × 2.00 × 100 mm). For the heat resistance study capillaries were filled using a drying chamber with a vacuum pump. Once the vacuum was achieved, it was broken and the rice solution rose through the capillaries, which were filled to a volume of 2/3 of their capacity. After that, the solution column was centred in the capillaries they were removed from the chamber and the open end was closed with a quick-drying silicone.

Before the heat resistance study spores were heat activated in order to create the conditions for them to germinate and grow in the culture medium. For the activation of

*B. cereus* spores the capillaries were placed in hooked racks designed for this type of study. The racks with the capillaries were immersed in a water bath (HAAKE N3) at 80 ° C ± 0.5 for 10 minutes.

Both the rice solution alone and the rice solution containing chitosan were heat treated at 90 95 100 and 105 ºC for different exposure times. A silicone oil bath (HAAKE DC5) was used for this treatment. For time cero (0) and for each treatment temperature. acapillary rack was removed after spore activation and was not heat-treated thus considered as control. The rest of the racks were withdrawn from the activation bath and immediately immersed in the oil bath at the selected temperature. A rack was removed at each time interval and immersed in ice water to stop the treatment.

Before the solution was plated the capillaries were cleaned with 96% ethanol and. using forceps the ends were split to extract the solution. The content of eight capillaries was deposited into sterile Eppendorf tubes. With the solution recovered from the capillaries two series of serial decimal dilutions (series A and B) were made up to 10-6 by duplicate. From each decimal solution 100 µL was plated in duplicate on nutrient agar (Scharlab, Barcelona, Spain) enriched with 1g/L starch (Scharlab, Barcelona, Spain) and incubated for 18-20 hours at 30ºC. Afterthe incubation time, a manual count of *B. cereus* colonies was carried out

### Statistical analysis

All statistical analyses including the one step nonlinear regression were performed using Statgraphics Centurion XVıSoftware (Addinsoft SARL. New York. NY. USA). Non-linear regression is a powerful technique for standardizing data analysis (Brown 2001), it allows obtaining the D and z values from survival curves at once

## Results

In the present work, the heat resistance of *Bacillus cereus* was studied in a rice substrate without insect chitosan and with insect chitosan at two concentrations.

The survival curves at each temperature tested in the study can be seen in Figures 1 to 4.

**Figure 1:**
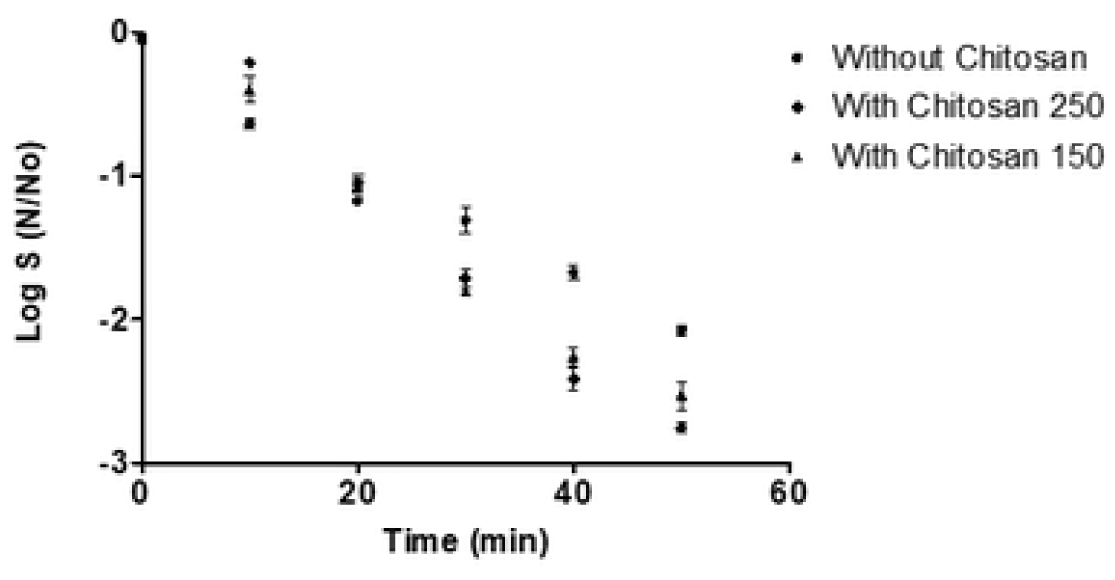
Survival curves for *Bacillus cereus* heated at 90 ºC

**Figure 2:**
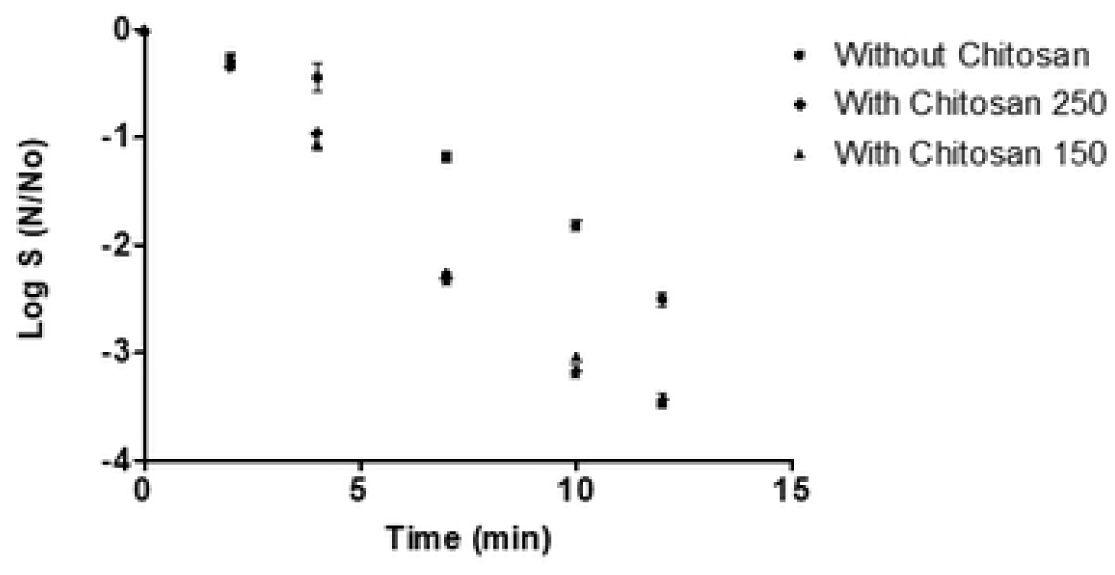
Survival curves for *Bacillus cereus* heated at 95 ºC

**Figure 3:**
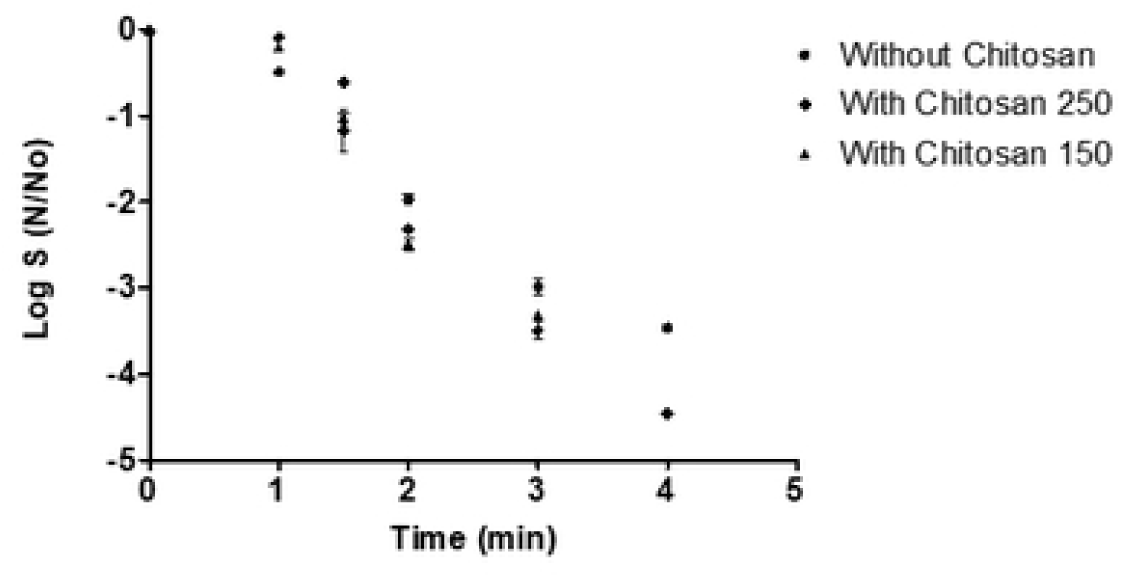
Survival curves for *Bacillus cereus* heated at 100 ºC

**Figure 4:**
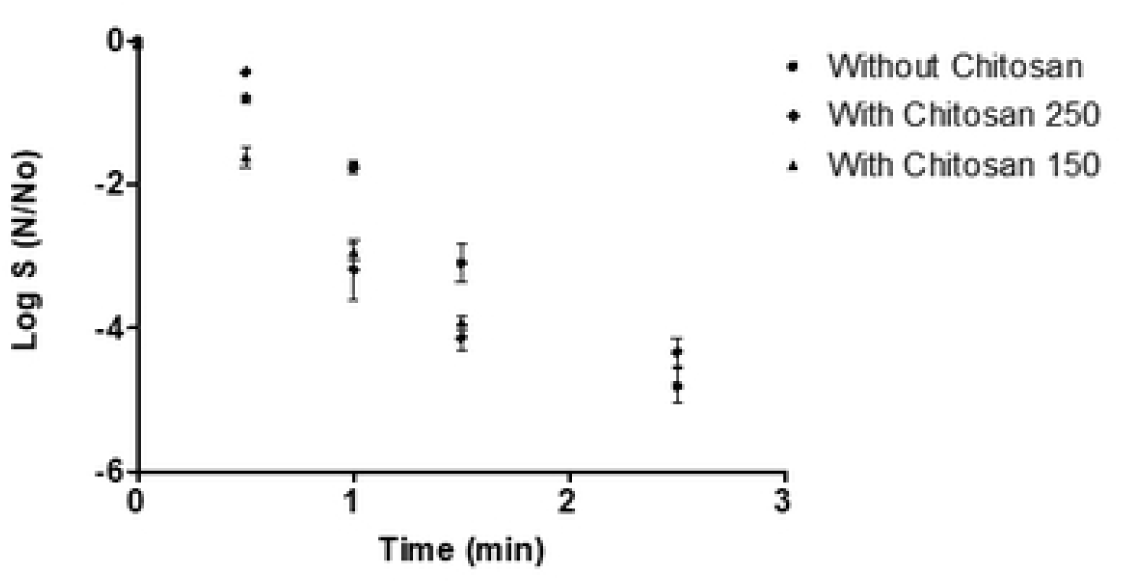
Survival curves for *Bacillus cereus* heated at 105 ºC

In general, at all temperatures studied. *B. cereus* spores were more resistant to heat in the rice substrate without chitosan. Regarding chitosan concentrations we also observed that for all temperatures the heat resistance of *B. cereus* spores was quite similar so the chitosan concentration in the heating medium did not affect the survival of these spores. The parameters defining the heat resistance of the spores were derived by a non-linear one-step fitting of the survival data. Nonlinear models often capture the relationships in a data set better than linear models. Perrin (2017) described the disadvantages of the usual linear least squares analysis of first- and second-order kinetic data and nonlinear least squares fitting was recommended as an alternative. In our study the value of the studentized residuals was in all cases two or less than two in any case three as absolute value this means that in no case the residuals exceed two standard deviations. Tables 1 2 and 3 show the estimation of the parameters that define the heat resistance of *B. cereus* spores D_T_ for each of the substrates and temperatures studied. Table 4 shows the z value for each of the studied substrate.

**Table 1:** Estimation of thermal resistance parameters by a nonlinear regression in a substrate without chitosan.

| Temperature (°C) | Estimated D value (min) | Standard Error Asymptotic |
| --- | --- | --- |
| 90 | 18.90 | 1.78 |
| 95 | 5.87 | 0.37 |
| 100 | 1.82 | 0.078 |
| 105 | 0.56 | 0.029 |

**Table 2:** Estimation of thermal resistance parameters by a nonlinear regression in substrate with chitosan 150 µg/mL.

| Temperature (°C) | Estimated D value (min) | Standard Error Asymptotic |
| --- | --- | --- |
| 90 | 15.47 | 0.97 |
| 95 | 4.27 | 0.18 |
| 100 | 1.18 | 0.050 |
| 105 | 0.32 | 0.023 |

**Table 3:** Estimation of thermal resistance parameters by a nonlinear regression in substrate with chitosan 250 µg/mL.

| Temperature (°C) | Estimated D value (min) | Standard Error Asymptotic |
| --- | --- | --- |
| 90 | 14.17 | 1.11 |
| 95 | 4.83 | 0.26 |
| 100 | 1.64 | 0.06 |
| 105 | 0.56 | 0.032 |

**Table 4:**
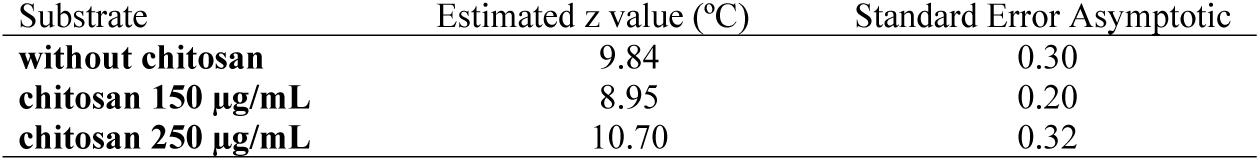
Estimated z values (ºC) in different substrates.

| Substrate | Estimated z value (°C) | Standard Error Asymptotic |
| --- | --- | --- |
| without chitosan | 9.84 | 0.30 |
| chitosan 150 µg/mL | 8.95 | 0.20 |
| chitosan 250 µg/mL | 10.70 | 0.32 |

The value of the parameter D_T_ estimated by the model is clearly higher in the substrate without chitosan than in the substrate containing chitosan which indicates greater spore resistance to heat treatment without chitosan as previously shown by the survival curves. Regarding the value of the parameter, D_T_ estimated by the model when chitosan is present little difference was found between the two chitosan concentrations. It seems that the effect of chitosan on the heat resistance of *B. cereus* spores does not depend on the concentration but rather on the molecular structure of chitosan and its interaction with elements of the bacterial spore during heating. With respect to the value of the z parameter estimated by the model varied between 8.9 and 10.7, those are quite common values for this microorganism (Fernandez et al. 1999, Alvarenga et al. 2018)

## Discussion

*Bacillus cereus* is a ubiquitous microorganism that can cause serious food safety issues especially in rice products and their derivatives. Proper characterization of its thermal resistance is essential for the design and development of suitable cooking processes. Likewise the prospect of using combined processes in this case with natural antimicrobials can pave the way to improving the safety of these widely consumed products around the world. Currently there is information on the effect of temperature and on the effect of chitosan separately on *B. cereus* spores.

Several works have reported the variation in the D_T_ and z values of the microorganism in different heating substrates. Pendurkaa and Kulkarnı(1989) studied the heat resistance of the spores of five *Bacillus* species including *B. cereus* in distilled water and pasteurized skim milk. The authors found that in all cases the spores survived the cooking conditions applied to the rice. At 100 ºC a D_T_ value of 19 min was shown by *B. cereus* in distilled water while *B. cereus* spores were completely inactivated in skim milk at the same temperature (100 ºC). This result indicates low levels of heat resistance. In the present work at 100 ºC a D_T_ value of 1.82 min was recorded when the spores were heated in a rice solution. However, the great variability that exists between *B. cereus* spores in relation to heat resistance is well known. Fernandez et al. (1999) studied the heat resistance of two *Bacillus cereus* strains isolated from cooked chilled foods containing vegetables and found D_T_ values between 0.22 and 2.5 min at 100ºC. More recently. Salwa Abu El-Nour. Ali Hammad (2013) found D_85_-values of *B. cereus* spores ranging from 24.9 to 35.2 min. D_90_-values ranging from 7.6 to 11.6 min. whereas D_95_-values ranged from 2.4 to 4.7 min. depending on the type of substrate. The values obtained in the present work are slightly higher probably due to the strain and substrate differences.

Regarding the z value. Fernandez et al. (1999) reported values of 8.1 and 8.4 ºC depending on the strain considered obtained on a reference substrate. Salwa Abu El-Nour. Ali Hammad (2013) reported z values of *B. cereus* spores suspended in different media ranging from 9.81 to 11.24 ºC. In the present work, the z value ranged from 8.9 ºC to 10.7 ºC depending on the substrate used. The z values obtained in the present work are in accordance with previously reported results; therefore these results can be considered a suitable reference to develop suitable cooking conditions for rice.

Today, chitosan is extensively studied given the multiple applications that it can have in both the food and the pharmaceutical industries. One of these applications is its use as a natural antimicrobial in food preservation. Ke et al. (2021) indicated that the broad-spectrum antimicrobial activity of chitosan offers great commercial potential for this product. Some studies have been published in which the effectiveness of chitosan against *B. cereus* has been demonstrated. Fernandes et al. (2009) found a relationship between the molecular weight of chitosan and its antimicrobial activity for both vegetative cells and spores of *B. cereus*. Mellegård et al. (2011) studied the inhibition of *B. cereus* spore outgrowth and multiplication by chitosan; they found chitosan exerts antimicrobial activity that appears to be concentration-dependent and related to the average molecular weight and fraction of acetylation of the chitosan used as antimicrobial.

Currently the industry is looking into combined treatments in which the different control measures are administered with lower intensities than when applied individually. In this way, pathogenic microorganisms are inactivated in a way that improves both the nutritional and sensory quality of food. In some cases, this combination is interesting because it can provide greater inactivation by heat than when heat is administered alone. There are no studies in the literature reporting the combination of heat treatment and chitosan to achieve control and inactivation of *B. cereus* in rice-based substrates. However, the effect of combining heat treatment or other control measures with natural antimicrobials has been reported in the scientific literature. Ueckert et al. (1998) reported that exposure to heat and nisin caused synergistic reductions of *Lactobacillus plantarum* viability. Huertas et al.. (2014) studied the combined effect of natural antimicrobials (nisin. citral and limonene) and thermal treatments on *Alicyclobacillus acidoterrestris* spores. Authors concluded that the antimicrobial agents tested did not affect the heat resistance of the spores; however, the antimicrobials were effective in controlling the growth of the microorganisms after the heat treatment. Kamdem et al. (2015) studied the effect of mild heat treatments on the antimicrobial activity of some essential oils. Authors indicated that the combination of temperature and those essential oils reduced the treatment time needed to inactivate 7 log cfu/mL of S*almonella* enteritidis. In the present work. a joint effect of heat and chitosan on *B. cereus* spores heat resistance was found, D_T_ values were in general lower on samples containing chitosan than in the sample without chitosan. Probably, the additive effect during heat treatment depends on the type of microorganism or the type of antimicrobial. In the present work, we found that the effect on D_T_ values was not dependent on chitosan concentration. It is possible that at this level the chitosan concentration does not play an important role but rather it is the molecular structure of the chitosan that facilitates the action of heat on the bacterial spores thus reducing the number of spores capable of germinating and growing.

## Conclusions

This study investigated the nature of the inactivation of *Bacillus cereus* spores by combining insect chitosan with heat treatment. The results indicated that the presence of chitosan regardless of its concentration produced reductions in the D_T_ value of *B. cereus* spores in a rice substrate. These findings pave the way to a better control of *B. cereus* during and after the cooking processes of rice and its derivatives making the combination of chitosan with heat treatment feasible in order to improve the safety of these types of products. These results also indicate that insect chitosan could be an alternative to crustacean chitosan, as antimicrobial in combination with heat treatment.

## Aknowledgements

We want to thank TRACE-RICE project. Reference Number AMD-1934-1, Horizon 2020, for the support for this Research,

